# SSCC: a novel computational framework for rapid and accurate clustering large single cell RNA-seq data

**DOI:** 10.1101/344242

**Authors:** Xianwen Ren, Liangtao Zheng, Zemin Zhang

**Author notes:** Emails: XR.

## Abstract

Clustering is a prevalent analytical means to analyze single cell RNA sequencing data but the rapidly expanding data volume can make this process computational challenging. New methods for both accurate and efficient clustering are of pressing needs. Here we proposed a new clustering framework based on random projection and feature construction for large scale single-cell RNA sequencing data, which greatly improves clustering accuracy, robustness and computational efficacy for various state-of-the-art algorithms benchmarked on multiple real datasets. On a dataset with 68,578 human blood cells, our method reached 20% improvements for clustering accuracy and 50-fold acceleration but only consumed 66% memory usage compared to the widely-used software package SC3. Compared to *k*-means, the accuracy improvement can reach 3-fold depending on the concrete dataset. An R implementation of the framework is available from https://github.com/Japrin/sscClust.

## 1. Introduction

Single cell RNA sequencing (scRNA-seq) has revolutionized transcriptomic studies by revealing the heterogeneity of individual cells with high resolution [1#x2013;6]. Clustering has become a routine analytical means to identify cell types, depict their functional states and infer potential cellular dynamics [4#x2013;10]. Multiple clustering algorithms have been developed, including Seurat [11], SC3 [12], SIMLR [13], ZIFA [14], CIDR [15], SNN-Cliq[16] and Corr [17], etc. These algorithms improve the clustering accuracy of scRNA-seq data greatly but often have high computational complexity, impeding the extension of these elegant algorithms to large-scale scRNA-seq datasets. With the rapid development of scRNA-seq technologies, the throughput has increased from initially hundreds of cells to tens of thousands of cells in one run [18]. Integrative analyses of scRNA-seq datasets from multiple runs or even across multiple studies further exacerbate the computational difficulties. Thus, algorithms that can cluster single cells both efficiently and accurately are needed.

To handle multiple large-scale scRNA-seq datasets, *ad hoc* computational strategies have been proposed by downsampling or convoluting large datasets to small ones [12, 19#x2013;21] or by accelerating the computation with new software implementation [22]. Such strategies have reached variable levels of success but have not adequately addressed the challenges. Considering the importance of efficient and accurate clustering tools for analyses of large-scale scRNA-seq data, here we propose a new computational framework based on machine learning techniques including feature engineering and random projection to achieve both improved clustering accuracy and efficacy. Benchmarking on various scRNA-seq datasets suggested that our new computational framework can reduce the computational complexity from O(n^2^) to O(n) while maintaining high clustering accuracy compared to the current solutions. Flexibility of the new computational framework allows our methods to be further extended and adapted to a wide range of applications for scRNA-seq data analysis.

## 2. Methods

### 2.1. Overview of the new computational framework

Among the available solutions to handle large scRNA-seq datasets, clustering with subsampling and classification [12, 19] has linear complexity, i.e. O(n). This framework generally consists of four steps (Figure 1a). First, a gene expression matrix is constructed by data preprocessing techniques including gene and cell filtration and normalization. Second, cells are divided into two subsets for clustering and classification separately by subsampling. Third, the subseted cells for clustering are grouped into clusters by *k*-means [23], hierarchical clustering [24], density clustering [25] or algorithms developed specially for scRNA-seq. Fourth, supervised algorithms such as *k*-nearest neighbors [26], support vector machines [27] or random forests [28] are used to predict the labels of other cells based on the clustering results in the third step. For simplicity, we referred this existing framework as SCC (**S**ubsampling-**C**lustering-**C**lassification). Because clustering is time-consuming and memory-exhaustive, limiting this step to a small subset of cells through subsampling can greatly reduce the computational cost from O(n^2^) to O(n) by leveraging the efficiency of supervised algorithms. However, classifiers built on the original gene expression data of a small subset of cells may be flawed and biased due to noise of the raw data and small number of cells, thus impairing the accuracy of label assignment for the total cells.

**Figure 1.**
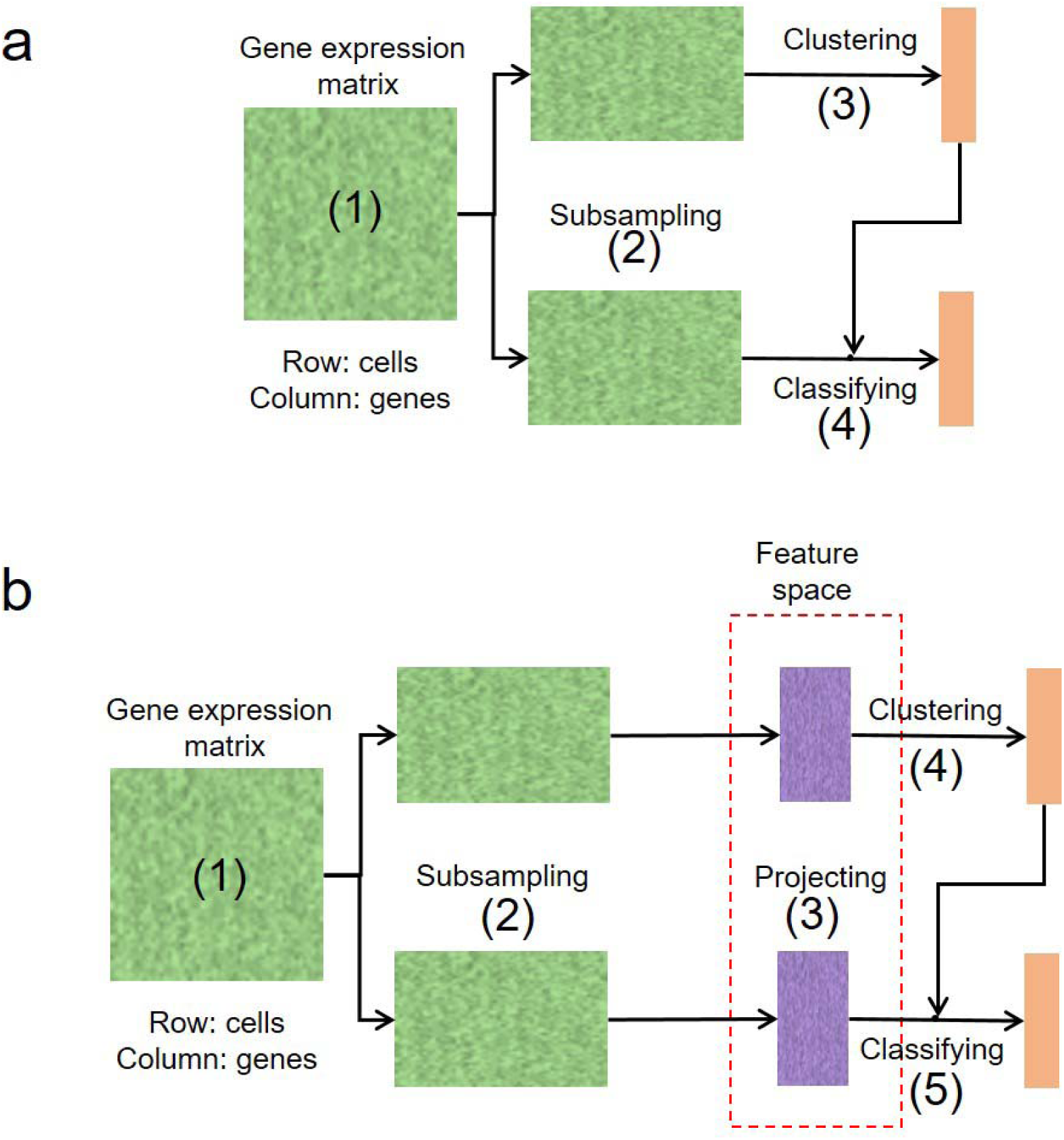
Two computational frameworks for rapid clustering large-scale single-cell RNA-seq datasets. a, the original computational framework proposed in SC3 to handle large single-cell RNA-seq datasets, which consists of four main steps: (1) constructing the gene expression matrix; (2) dividing the matrix into two parts through cell subsampling; (3) clustering the subsampled cells; (4) classifying the unsampled cells into clusters formed by the subsampled cells. b, the new computational framework we proposed to handle large single-cell RNA-seq datasets. Different from the original framework, a feature construction step is added before clustering and classifying. Thus the whole framework is composed by five steps: (1) constructing the gene expression matrix; (2) dividing the matrix into two parts through cell subsampling; (3) projecting those subsampled/unsampled cells into feature space; (4) clustering the subsampled cells in the feature space; (5) classifying the unsampled cells into clusters formed by the subsampled cells in the feature space.

Here we proposed a new computational framework for clustering large scRNA-seq data by adding a feature engineering/projecting step into SCC (Figure 1b). Similar to SCC, a gene expression matrix is first constructed through gene and cell filtrations and normalization (Step 1 in Figure 1b), and is then split randomly into two subsets for clustering and classification separately (Step 2 in Figure 1b). Unlike SCC which directly use the raw data of gene expression, our framework projects cells into a feature space (Step 3 in Figure 1b) for clustering (Step 4 in Figure 1b) and classification (Step 5 in Figure 1b). As the new framework is characterized by a Subsampling-Featuring-Clustering-Classification strategy, we named it as SFCC. Specifically, we divide feature construction into two steps. First, feature extraction techniques are applied to cells subject to clustering. Then, according to the selection of feature extraction methods, cells for classification are projected into the built feature space. Many established techniques in the machine learning field can be exploited in these two steps. For example, principal component analysis (PCA) [29] can be used to first construct features for cells undergoing clustering while the resultant loading vectors can be used as linear transformations to project cells for classification into the feature space. Selecting different algorithms in each step of the SFCC framework will then form different pipelines for clustering large-scale scRNA-seq datasets. To reduce the total number of algorithmic combinations, here we focus on comparing the performance between various feature engineering algorithms. We hold algorithms for gene and cell filtration, normalization, subsampling and classification as the algorithms frequently used in practice (see below for details). The existing SCC strategy can be treated as a special case of SFCC in which the original data space is the feature space.

Feature engineering techniques involved in this study include distance-based methods (Euclidean and cosine), correlation-based methods (Pearson [30] and Spearman [31] correlations) and a neural network-based method (Autoencoder) [32]. For distance and correlation based methods, the distance/correlation matrix for cells subject to clustering was directly used as their features, and the distance/correlation matrix between cells subject to classification and clustering were used to construct features for cells undergoing classification. For Autoencoder, the gene expression data of cells for clustering were used to train a neural network model first and then all cells were projected into a feature space through the encoding function of the trained model. To obtain evaluation results independent of clustering algorithms, we used silhouette values [33] to examine the global performance of these feature engineering methods. Upon the global evaluation, we then select the most effective method (SSCC: SFCC with Spearman correlation as the feature construction method) to do further evaluations.

### 2.2. scRNA-seq datasets used in this study

We used seven scRNA-seq datasets to evaluate the effects of clustering in feature space. Detailed descriptions of all the seven datasets are listed below.

1. The Kolodziejczyk dataset [34]. This dataset contains 704 cells with three clusters that were obtained from mouse embryonic stem cells with different culture conditions. About 10,000 genes were profiled with high sequencing depth (average 9 million reads per cell) by using the Fluidigm C1 system and applying the SMARTer Kit to obtain cDNA and the Nextera XT Kit for Illumina library preparation.
2. The Pollen dataset [8]. This dataset contains 249 cells with eleven clusters, which contain skin cells, pluripotent stem cells, blood cells, neural cells, etc. Either low or high sequencing depth based on the C1 Single-Cell Auto Prep Integrated Fluidic Circuit, the SMARTer Ultra Low RNA Kit and the Nextera XT DNA Sample Preparation Kit was used to depict the gene expression profiles of individual cells.
3. The Usoskin dataset [9]. This dataset contains 622 mouse neuronal cells with four clusters i.e., peptidergic nociceptors, nonpeptidergic nociceptors, neuroflament containing, and tyrosine hydroxylase containing. The neuronal cells were picked with a robotic cell-picking setup and positioned in wells of 96-well plates and then were subject to RNA-seq with 1.14 million reads per cell.
4. The Zeisel dataset [10]. This dataset contains 3,005 cells from the mouse brain with nine major subtypes. The gene expression levels were estimated by counting the number of unique molecular identifiers (UMIs) obtained by Drop-seq.
5. The Zheng dataset [5]. This dataset contains 5,063 T cells from five patients with hepatocellular carcinoma of which nine subtypes of samples were prepared according to the tissue types and cell types and then were subject to Smart-seq2 for profiling the gene expressions.
6. The PBMC 68k dataset [18]. Gene expression of 68,578 peripheral blood mononuclear cells (PBMC) of a healthy human were profiled by the 10X Genomics GemCode platform, with 3’ UMI counts used to quantify gene expression levels by their customized computational pipeline. This cell population includes eleven major immune cell types.
7. The Macoskco dataset [19]. This dataset contains 49,300 mouse retina cells without known distinct clusters. The gene expression levels were estimated by counting the number of UMIs obtained by Drop-seq, which were further clustered into 39 cell subtypes by the authors based on the Seurat algorithm.

### 2.3. Data preprocessing

The first four datasets (i.e., the Kolodziejczyk, Pollen, Usoskin and Zeisel datasets) has been widely used for evaluating clustering algorithms, of which the preprocessed data have been included in the SIMLR software package for testing use. We downloaded these four datasets from the Matlab subdirectory of the SIMLR package, and then selected the top 5000 most informative genes (with both the average and the standard deviation of log2-transformed expressing values more than 1) for subsequent analysis. If the gene number in the data was smaller than 5000, then all the genes were retained for further analysis. For the Zheng dataset, one patient (P0508) was selected for comparison of different clustering algorithms, which had 1,020 T cells with eight subtypes defined by the tissue sources and the cell surface markers. The same criteria were used to filter genes and then the transcripts per million (TPM) values were used to do clustering evaluations. For the PBMC 68k dataset, the preprocessing pipeline of the original paper [18] was used to prepare data for clustering (https://github.com/10XGenomics/single-cell-3prime-paper). For the Macoskco dataset, the UMI counts were used for evaluations without gene filter.

### 2.4. Consistency between true labels and the original and projected data

The silhouette value [33] is used to measure the consistency between the true labels and the original and projected data. Given a dataset with n samples and a clustering scheme, a silhouette value is calculated for each sample. For a sample *i*, its silhouette value *s_i_* is calculated according to the following formula:

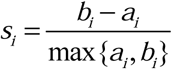

Where *a_i_* is the average dissimilarity of *i* to samples in its own cluster and *b_i_* is the lowest average dissimilarity of *i* to any other cluster of which *i* is not a member. *s_i_* is between −1 and 1. Closeness to 1 means it is well matched to its cluster while closeness to −1 means that *i* would be more appropriate if it was clustered in its neighboring cluster. For each feature construction method, the median silhouette value of the whole cells after projection was used to evaluate its consistency with the true cluster labels. The fraction of cells that have silhouette values increased after projection compared to the original data (i.e., the fraction of cells above the diagonal in Figure 2) was also used to evaluate the feature construction method.

**Figure 2.**
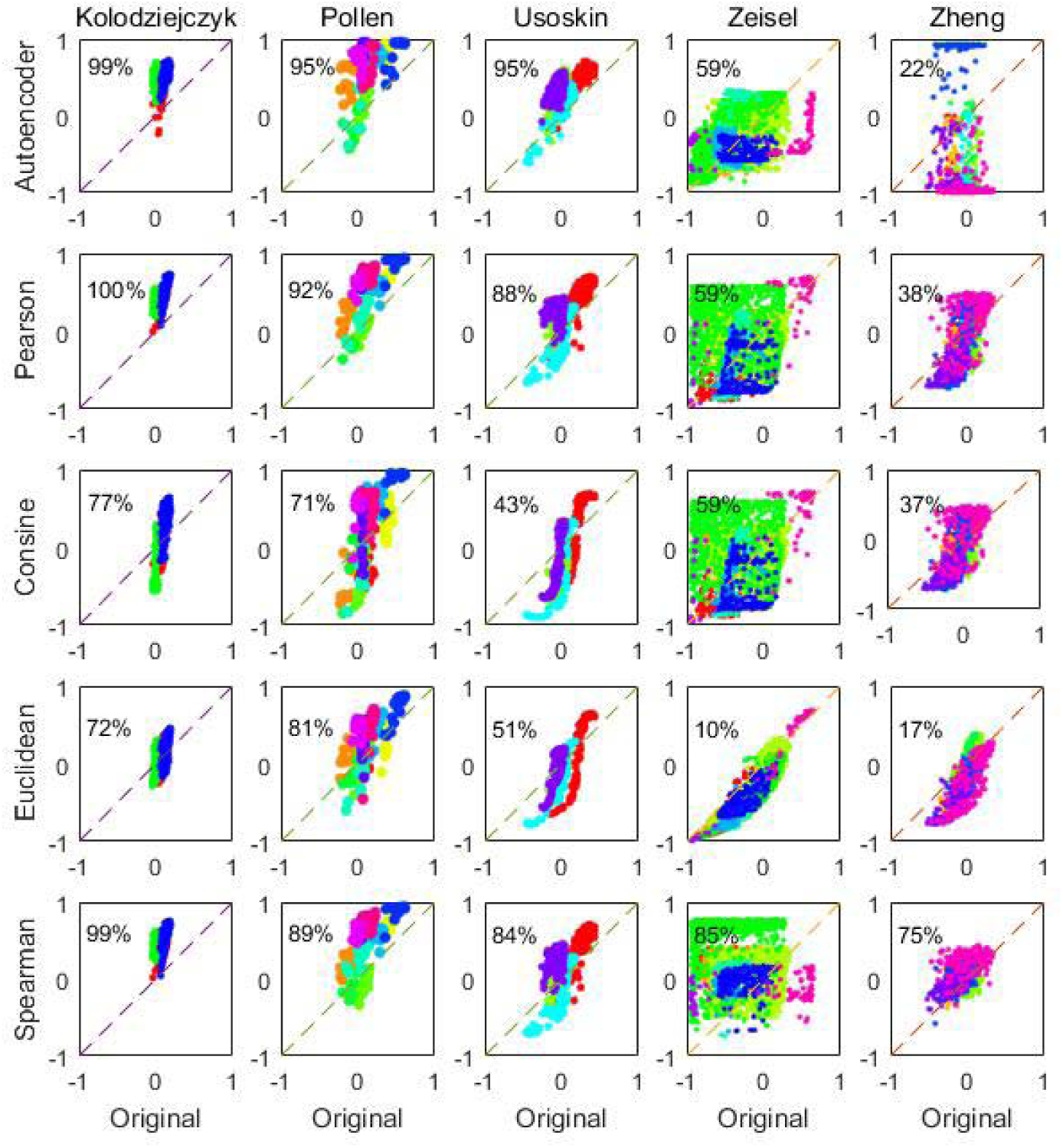
Features constructed by Spearman correlations consistently have better consistency with true cluster labels than the original data on all the five datasets (Kolodziejczyk, Pollen, Usoskin, Zeisel and Zheng). X axis: consistency between true cluster labels and the original data; Y axis: consistency between true cluster labels and various features extracted by Autoencoder, Pearson correlation, consine, Euclidean distance and Spearman correlation. Silhouette values6 were used to measure the consistencies and calculated for each cell by using the silhouette function of Matlab R2016a, with 1 representing perfect match while −1 indicating that the cell might be mis-clustered. The digital numbers in each plot suggest the fraction of cells whose clustering are improved in the feature space compared to those based on the original data (i.e., above the diagonals).

### 2.5. Clustering accuracy/consistency evaluation

Normalized mutual information (NMI)[35] was used to evaluate the accuracy of various clustering results. Given two clustering schemes 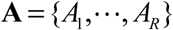 and 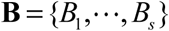, the overlap between **A** and **B** can be represented through the contingency table **C** (also named as confusion matrix) of size *R*×*S*, where *C_ij_* denotes the number of cells that clusters *A_i_* and *B_j_* share. The normalized mutual information *NMI*_(**A**,**B**)_ of the two clustering schemes **A** and **B** is defined as follows.

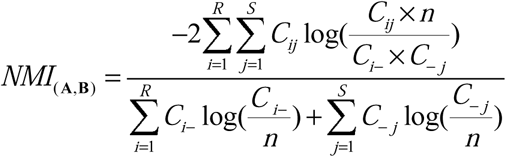

Where *n* is the number of total cells, *C_i_*_−_ is the number of cells assigned to cluster *i* in the clustering scheme **A** and *C*_−*j*_ is the number of cells assigned to cluster *j* in the clustering scheme **B**. If **A** is identical to **B**, *NMI*_(**A,B**)_ =1. If **A** and **B** are completely different, *NMI*_(**A,B**)_ =0. When true cluster labels were available, the NMI values between true cluster labels and various clustering results were used to evaluate the clustering accuracy. When true cluster labels were not available, NMI was used to evaluate clustering consistency between different subsampling rates in this study. Besides NMI, we also used rand index and adjusted rand index to evaluate clustering accuracy and consistency and obtained similar observations.

### 2.6. Clustering and classification algorithms

Many clustering algorithms are available. We selected five widely-used clustering algorithms in this study to evaluate the impacts of Spearman correlation-based feature construction method. These five algorithms include three general clustering algorithms which were designed initially not for scRNA-seq data (affinity propagation (AP)[36], *k*-means[23] and *k*-medoids[37]) and two algorithms that were specially designed for clustering of scRNA-seq data (SC3[12] and SIMLR[13]). *K*-means and *k*-medoids are pure algorithms that partition samples into groups while AP, SC3 and SIMLR inherently include feature construction techniques. All these clustering algorithms were evaluated on the five small-scale datasets while on the PBMC 68k dataset only SC3 was evaluated and on the Macoskco dataset only k-means was evaluated for simplicity. Parameters (ks = 10:12, gene_filter = FALSE, biology = FALSE, svm_max = 5000) were used for SC3 (default) while (ks = 11, gene_filter = FALSE, biology = FALSE, svm_max = 200) were used for SC3+SSCC. On the Macoskco dataset, ~5% and 10% cells were randomly picked out to do clustering. We used the *k*-nearest neighbors algorithm for classifying unsubsampled cells, which is robust to parameter selection.

## 3. Results

### 3.1 Feature construction can greatly improve the consistency of cell features and the reference cell labels

We calculated the silhouette values to evaluate the consistency between cell features extracted by various methods and the reference labels. Silhouette values are frequently used to indicate whether a sample is properly clustered. But here we can use silhouette values to reversely indicate whether the extracted features are properly consistent with the reference cell labels. By comparing with silhouette values of the original scRNA-seq data, we observed that most of the evaluated feature-extracting methods can improve the silhouette values for many cells on multiple datasets (Figure 2). On the Kolodziejczyk [34] and Pollen [8] datasets, all the five feature-extraction methods improved the silhouette values compared with the original data. On the Usoskin [9] and Zeisel [10] datasets, all methods showed good performance except Euclidean and cosine. On the Zheng [5] dataset, most methods failed except the Spearman correlation method. The Spearman correlation-based method consistently improved the accordance between cell features and labels on all the five datasets. Considering the robustness of Spearman correlation-based method and the great improvement of silhouette values of single cells, we evaluated this **S**pearman correlation-based **S**ingle-**C**ell **C**lustering framework (*abbr.* SSCC) in the next sections.

### 3.2 Clustering accuracy of the total cells is enhanced in feature space when subsampling is applied

We observed that the improvements of silhouette scores by SSCC were robust to subsampling fluctuations (Figure 3). On all five evaluated datasets, the silhouette values of Spearman correlation-based features were almost unchanged with subsampling rates (Figure 3), implying that features constructed upon SSCC with low subsampling rates may contain information approximate to those upon total cells.

**Figure 3.**
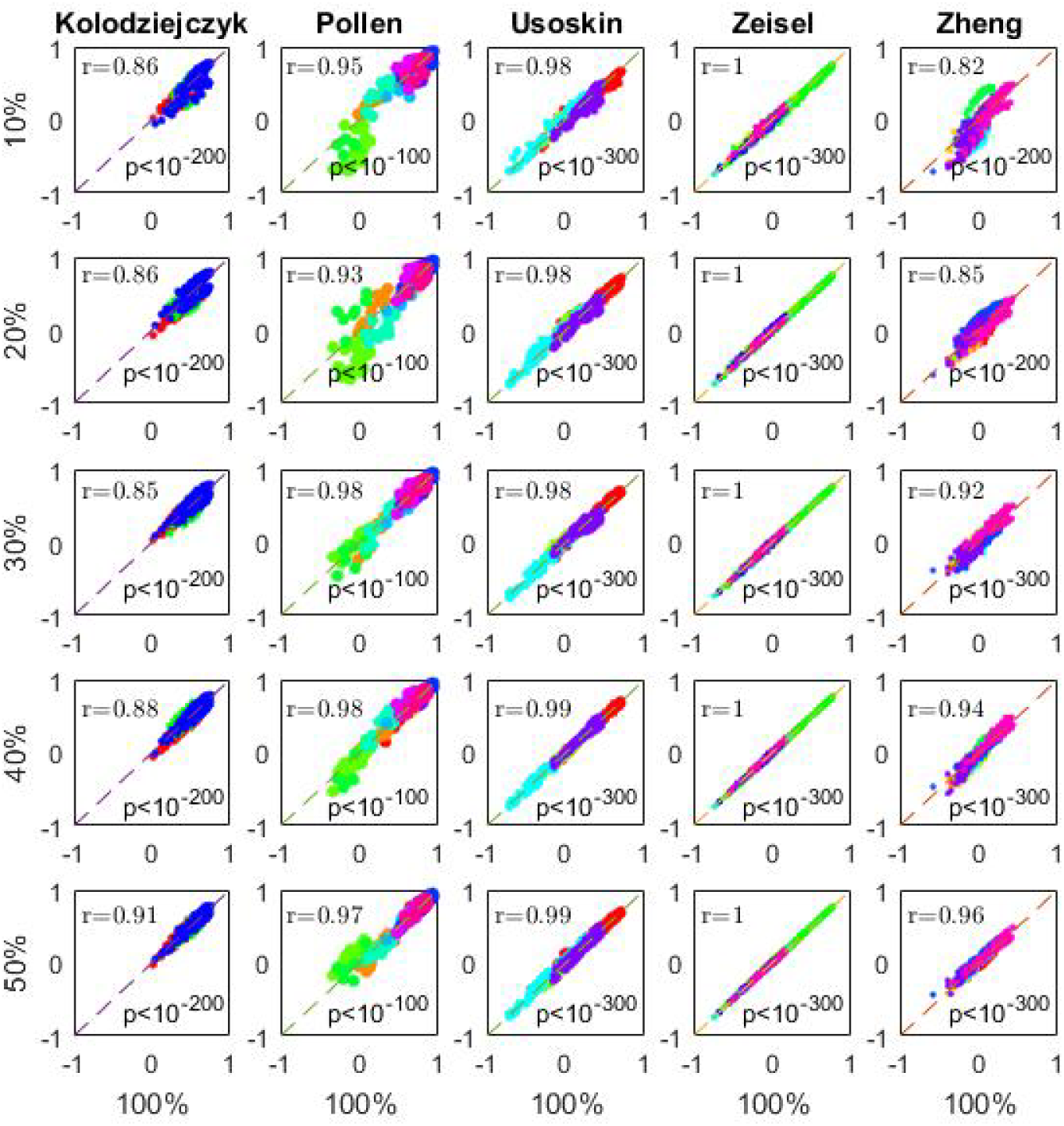
Silhouette values between Spearman correlation features and true cluster labels were independent on subsampling rates. X axis: silhouette values of Spearman correlation features constructed with 100% cells. Y axes: silhouette values of Spearman correlation features constructed with10%, 20%, 30%, 40% and 50% cells in each dataset. Pearson correlation was calculated to evaluate the consistency between subsampled and unsubsampled features, of which the correlation coefficients (r) were plotted in the upper triangles while the corresponding p values were plotted in the lower triangles. It is evident that silhouette values of features constructed upon subsampled cells were almost the same as those constructed upon all the cells in the datasets. The independence of such Spearman correlation-based features on subsampling rates will greatly reduce the computational complexity of feature construction (from O(n^2^) to O(n)), enabling clustering of large scRNA-seq datasets be finished with high efficacy.

We further evaluated whether the improved silhouette values can be translated into clustering accuracy. By evaluating five clustering algorithms including *k*-means, *k*-medoids, affinity propagation, SC3 and SIMLR, we observed that SSCC can significantly improve the clustering accuracy for all the five clustering algorithms on all the benchmark datasets (Figure 4). The accuracy improvements measured by ΔNMI can reach from 0.12 to 0.60 on the Kolodziejczyk dataset, 0.04 to 0.19 on the Pollen dataset, 0.14 to 0.37 on the Usoskin dataset, 0.02 to 0.28 on the Zeisel dataset, and 0.10 to 0.28 on the Zheng dataset, which depended on the specific selection of algorithms and subsampling rates. Other accuracy metrics including rand index, adjusted rand index and adjusted mutual information revealed the same trends, suggesting that SSCC can greatly enhance the clustering power of multiple algorithms when subsampling is used.

**Figure 4.**
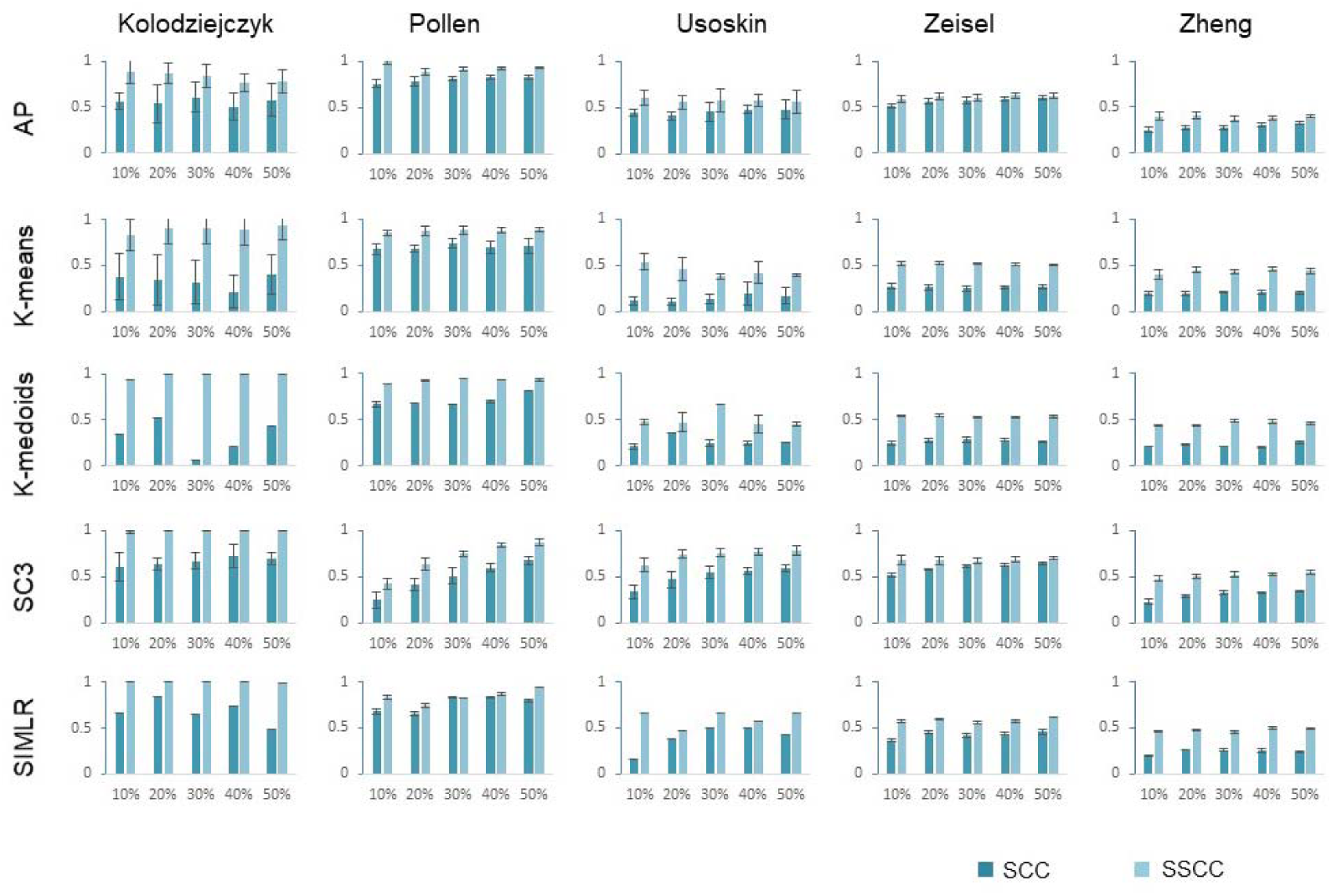
Performance comparison between SCC and SSCC. For each plot, the X axis was the subsampling rate, i.e., the percentage of cells used in clustering, and the Y axis was the clustering accuracy measured by NMI. For each subsampling rate, calculations were repeated for ten times, based on which the average and the standard deviation of the clustering accuracy were calculated and plotted. It is evident that SSCC significantly improved clustering accuracy for almost all algorithms, all scRNA-seq datasets and all subsampling rates.

### 3.3 Clustering consistency between different subsampling runs is also greatly improved with SSCC

In practice, the reference cell labels are generally unknown. The confidence of clustering results is often evaluated with the consistency between different algorithms. Because of subsampling fluctuations, clustering results based on SCC are inconsistent among different subsamplings. However, in the new framework of SSCC, the consistency was much improved for all evaluated clustering algorithms on all datasets (Figure 5). On the Kolodziejczyk dataset, all the five clustering algorithms reached consistency larger than 0.5 in SSCC while the corresponding consistency in SCC was much smaller. On the Pollen dataset, SSCC still showed better performance than SCC although both frameworks had high clustering consistency. Similar trends were observed on the Usoskin, Zeisel and Zheng datasets.

**Figure 5.**
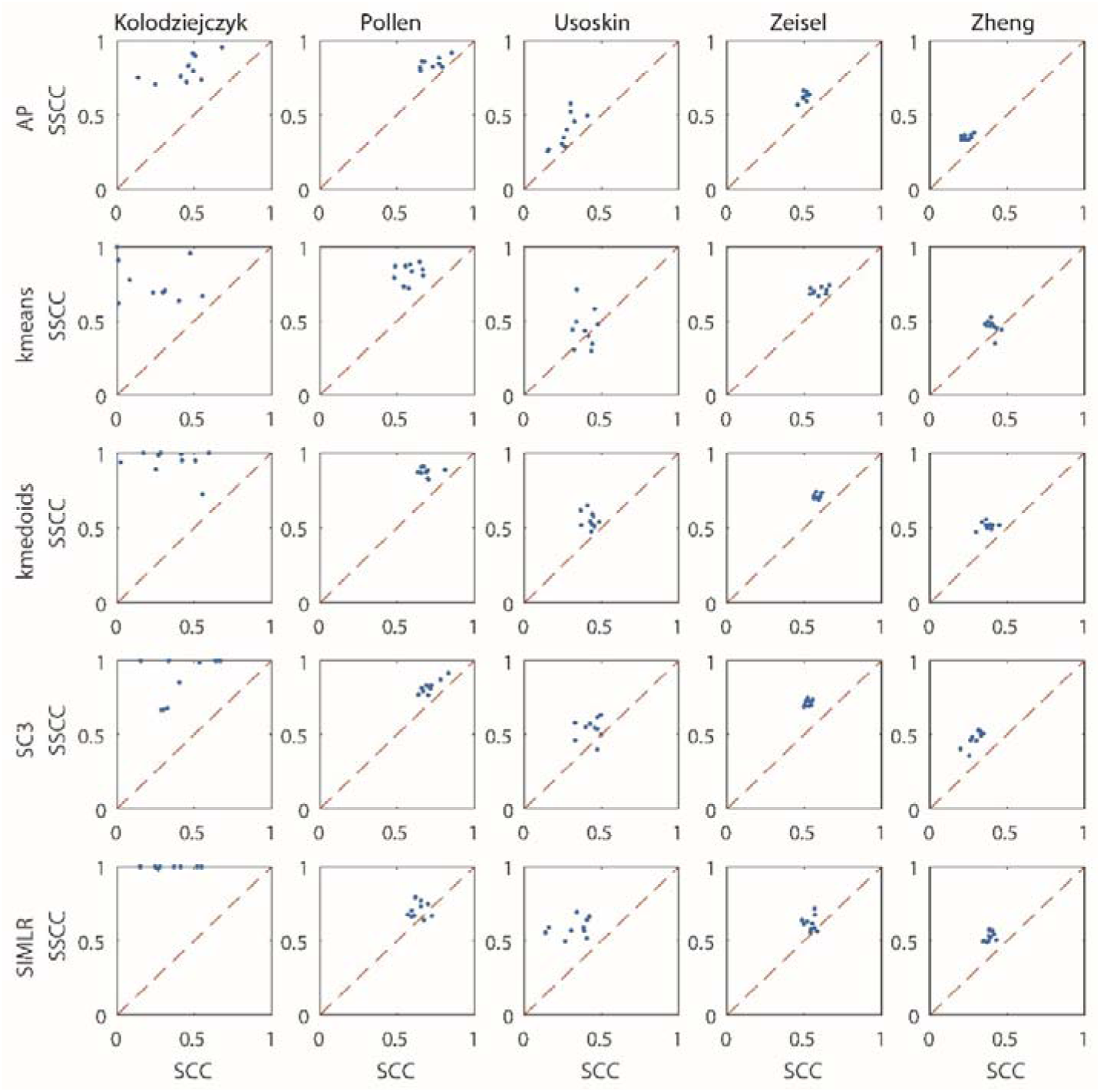
Comparison of clustering consistency between SSCC and SCC. X axis: consistency (measured by NMI) of clustering upon 10% cells to that based on 50% cells with SCC. Y axis: consistency (measured by NMI) of clustering upon 10% cells to that based on 50% cells with SSCC. Subsamplings were repeated for ten times and each subsampling result was processed by five clustering algorithms (AP, k-means, k-medoids, SC3, and SIMLR).

### 3.4 Application of SSCC to large scRNA-seq datasets with or without reference cell labels

Besides the former five scRNA-seq datasets, we further tested SSCC on two additional large scRNA-seq datasets. One is the PBMC 68k dataset [18], which contains 10X Genomics-based expression data for 68,578 blood cells from a healthy donor. The other is the Macoskco dataset [19] containing 49,300 mouse retina cells but without cell labels determined by experimental methods. The large cell numbers generally prohibit classic scRNA-seq clustering algorithms running on a desktop computer, and thus provide two realistic examples to demonstrate the performance of SCC and SSCC.

On the PBMC 68k dataset, we compared SSCC with SCC by using SC3 [12] as the clustering algorithm. The SC3 software package inherently applied an SCC strategy to handle large scRNA-seq datasets. By default, if a dataset has more than 5,000 cells, the SCC strategy will be triggered, with 5,000 cells randomly subsampled for SC3 clustering and the other cells for classification by SVM. We applied SC3 to the PBMC 68k dataset on a desktop computer with 8GB memory and 3GHz 4-core CPU and repeated ten times. The average clustering accuracy of SC3 was 0.48 measured by NMI, the calculation lasted 99 minutes on average, and the maximum memory usage exceeded 5.6 GB (Figure 6a). With our SSCC strategy, the average clustering accuracy reached 0.59, ~21% increase over SC3 (default). The computation time was dramatically reduced to 2.2 minutes on average, reaching 50-fold acceleration. The maximum memory usage of SC3+SSCC was 3.7GB, saving >33% compared to that of SC3 (default).

**Figure 6.**
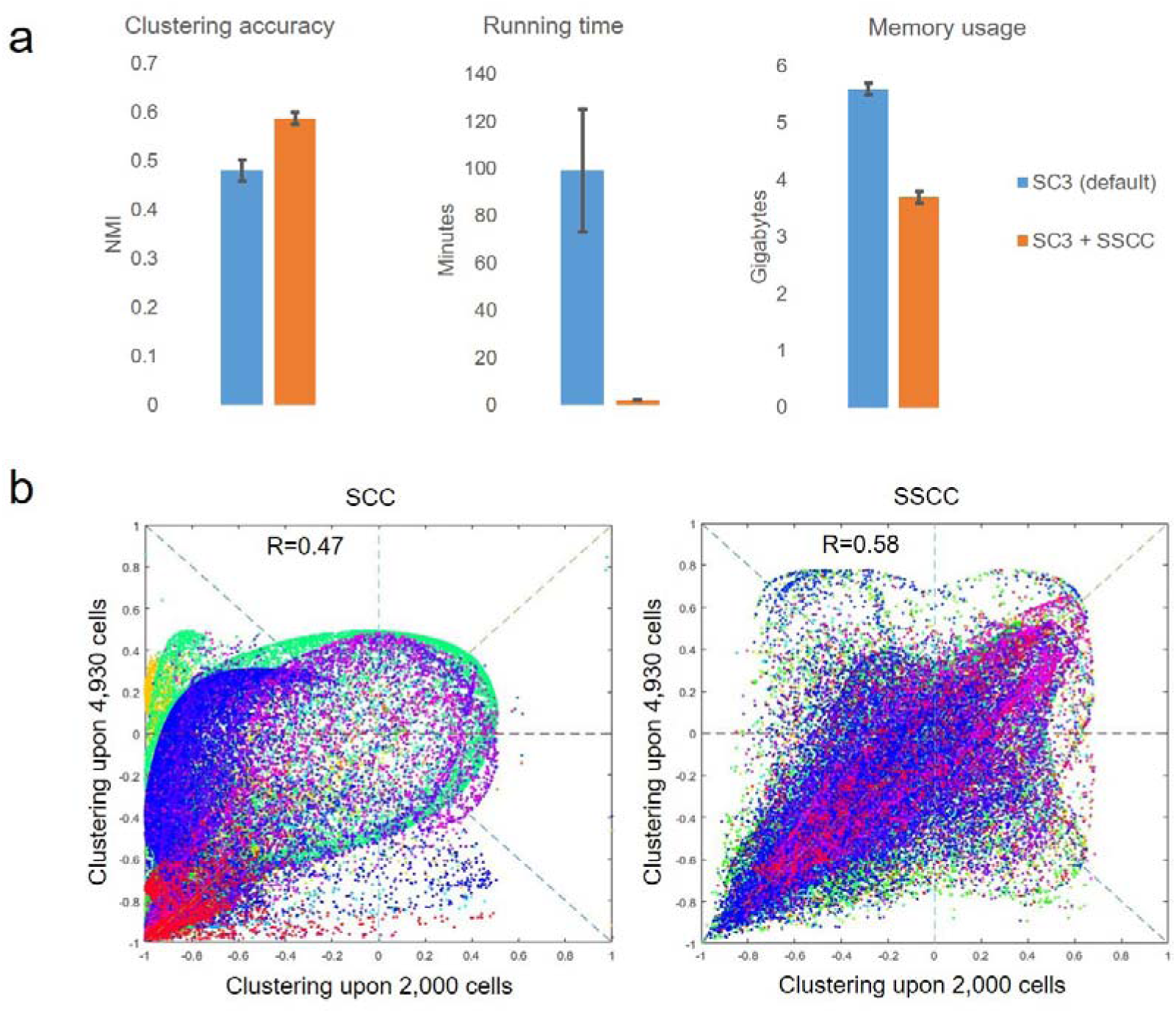
Performance evaluation of SSCC on two extremely large scRNA-seq datasets. a, Performance comparison between SC3 (default) and SC3+SSCC on the PBMC 68k dataset. SC3 (default) inherently applied the SCC strategy to deal with large scRNA-seq datasets, of which 5,000 cells were subsampled by default for clustering and other cells were subject to classification based on SVM. For SC3+SSCC, an extremely low number of cells (n=200) were subsampled to construct cell features and then for clustering and classification. b, Consistency comparison between SSCC and SCC on a large scRNA-seq dataset with 49,300 mouse retina cells. The silhouette values of two clustering schemes (by 2000 and 4930 cells separately) with the SCC framework (the left panel) were small and relatively weak-correlated to each other (R=0.47, Pearson’s correlation). The silhouette values of two clustering schemes (by 2000 and 4930 cells separately) with the SSCC framework were large and highly correlated to each other (R=0.58, Pearson’s correlation). Cells were colored according to cluster labels based on ~10% cells and original expression data.

On the Macoskco dataset, with *k*-means for clustering and *k*-nearest neighbors for classification, the SCC strategy resulted great average silhouette difference (−0.80 − −0.51=0.29) between two subsampling schemes (upon 5% and 10% cells) while the difference with SSCC became negligible. The NMI value between the two clustering results with SCC was 0.60 while that with SSCC was 0.69. Pearson correlations of silhouette values between the two clustering schemes were increased from 0.47 to 0.58 when switching from SCC to SSCC (Figure 6b). All these metrics suggest that SSCC can not only greatly improve the clustering efficiency and accuracy for large-scale scRNA-seq datasets, but also can greatly improve the consistency.

## 4. Discussion

The availability of large-scale scRNA-seq data raise urgent need for efficient and accurate clustering tools. Currently a few scRNA-seq analysis packages have been proposed to address this challenge. Of these tools, SC3 [12], Seurat [11] and dropClust [20] adopted a subsampling-clustering-and-classification strategy, bigScale [21] employed a convolution strategy to merge similar single cells into mega-cells by a greedy-searching algorithm, and SCANPY used Python as the programming language to accelerate the clustering process. Although these strategies greatly boost the efficiency of large scRNA-seq data analysis, much room still exist for further improvement. Particularly the subsampling-clustering-and-classification strategy suffers from biases introduced by subsampling which may greatly decrease the clustering accuracy and robustness, although it can reduce the computational complexity from O(n^2^) to O(n). Here we introduced feature engineering and projecting techniques into the SCC framework and proposed SFCC as an alternative. Specially, with Spearman correlations as the feature engineering and projecting methods, we formulated a framework named as SSCC, which can significantly improve clustering accuracy and consistency for many general and specially designed clustering algorithms. Evaluations on real scRNA-seq datasets, which covers a wide range of scRNA-seq technologies, sequencing depth and organisms, demonstrated the robustness of the superior performance of SSCC. Therefore, SSCC is expected to be a useful computational framework that can further unleash the great power of scRNA-seq in the future.

## AVAILABILITY

The five scRNA-seq datasets involved in this study and the Matlab codes are available from https://sites.google.com/site/renxwise/home/sccs. The newly proposed clustering framework is implemented as an R package available from https://github.com/Japrin/sscClust.

